# A targeted multi-omic analysis approach measures protein expression and low abundance transcripts on the single cell level

**DOI:** 10.1101/700534

**Authors:** Florian Mair, Jami R. Erickson, Valentin Voillet, Yannick Simoni, Timothy Bi, Aaron J. Tyznik, Jody Martin, Raphael Gottardo, Evan W. Newell, Martin Prlic

## Abstract

High throughput single-cell RNA sequencing (sc-RNAseq) has become a frequently used tool to assess immune cell function and heterogeneity. Recently, the combined measurement of RNA and protein expression by sequencing was developed, which is commonly known as CITE-Seq. Acquisition of protein expression data along with transcriptome data resolves some of the limitations inherent to only assessing transcript, but also nearly doubles the sequencing read depth required per single cell. Furthermore, there is still a paucity of analysis tools to visualize combined transcript-protein datasets.

Here, we describe a novel targeted transcriptomics approach that combines analysis of over 400 genes with simultaneous measurement of over 40 proteins on more than 25,000 cells. This targeted approach requires only about 1/10 of the read depth compared to a whole transcriptome approach while retaining high sensitivity for low abundance transcripts. To analyze these multi-omic transcript-protein datasets, we adapted One-SENSE for intuitive visualization of the relationship of proteins and transcripts on a single-cell level.

## Introduction

While pioneering work almost 20 years ago illustrated the ability to study the transcriptome at the single-cell level (Chiang and Melton, 2003; Phillips and Eberwine, 1996), recent advances in microfluidics and reagents allow the high-throughput analysis of transcripts of 10^4^ single cells in one experiment (Jaitin et al., 2014; Klein et al., 2015; Macosko et al., 2015). Although several methods have been developed for this purpose, currently the most widely adopted platform is a droplet-based microfluidic system commercialized by 10x Genomics (Zheng et al., 2017).

Though analysis of transcript expression on the single cell level is a powerful tool to characterize the relationship and functional properties of cells, it is imperative to consider the relationship between transcript and protein when trying to extrapolate biology. Typically, transcripts are expressed at a much lower level than proteins – for example, murine liver cells have a median copy number of 43,100 protein molecules but only 3.7 RNA molecules per gene (Azimifar et al., 2014). Similarly, the dynamic range of expression is much greater for proteins with copy numbers spanning about 6-7 orders of magnitude while transcript copy numbers span about 2 orders of magnitude (Schwanhausser et al., 2011). Finally, the correlation of gene expression and protein expression has been estimated to have a Pearson correlation coefficient between 0.4 (Schwanhausser et al., 2011) and 0.6 (Azimifar et al., 2014). These discrepancies in transcript and protein expression patterns are relevant for the biological interpretation of single cell transcriptome data, but also pose analytical challenges. Suitable approaches are required to visualize the data despite the pronounced differences in abundance and dynamic range of expression.

The parallel measurement of transcript and protein phenotype by sequencing has been recently reported as cellular indexing of transcriptomes and epitopes (CITE-seq) (Stoeckius et al., 2017) or RNA expression and protein sequencing (REAP-seq) (Peterson et al., 2017). These technologies leverage existing sc-RNAseq platforms that use an unbiased whole transcriptome (WTA) detection approach capturing cellular mRNA via its poly-A tail, and utilize oligonucleotide-labelled antibodies (carrying unique barcodes and also a poly-A tail) to interrogate surface protein abundance. Typically, current droplet-based WTA approaches result in the detection of ~1000 unique transcripts per single cell for the transcriptome (with a substantial fraction of these transcripts encoding ribosomal proteins), while antibody panels of up to 80 targets have been reported (Peterson et al., 2017).

Though proof-of-principle for this technology has been established, it remains unclear how the sequencing-based antibody detection compares to established flow cytometry-based assays in different experimental settings with regards to capturing the dynamic range of protein expression and identifying low abundance protein expression. In addition, the combined WTA plus protein approach can quickly become resource intensive. Finally, droplet-based WTA pipelines may still miss specific transcripts of interest if they are below the limit of detection, with current high throughput chemistries capturing an estimated 10% of the total cellular mRNA (Zheng et al., 2017).

Here, we report using a high throughput (>10^4^ single cells) targeted transcriptomic approach employing nanowells to capture single cells (Rhapsody platform, commercialized by BD Biosciences) (Fan et al., 2015) in combination with oligonucleotide-barcoded antibodies (termed AbSeq). Specifically, we simultaneously interrogated over 400 immune-related genes and 41 surface proteins that are commonly used for immunophenotyping. We found that this targeted approach was efficient at detecting low-abundance transcripts while only requiring about 1/10 of the sequencing read depth needed for WTA, indicating that targeted transcriptomics is a sensitive and cost-efficient alternative when the focus is on interrogating defined transcripts. Of note, this approach clearly separated different memory T cell subsets as well as regulatory T cells (Tregs) solely based on transcript information, which is often difficult due to the low amount of RNA recovered from T lymphocytes (Zheng et al., 2017). Furthermore, we used 30-parameter fluorescent-based flow cytometry to measure the same proteins targets as in the multi-omic assay. Our data indicate that the validation of oligonucleotide-barcoded antibody panels is necessary for meaningful interpretation of the multi-omic data.

To demonstrate the sensitivity and robustness of the system we analyzed T and NK cells before and after one hour of stimulation, revealing an unexpected disconnect in transcript and surface expression levels of the commonly used early activation marker CD69. Analysis of chemokine expression showed distinct phenotypes within the CD8^+^ T cell population as early as 60 minutes after stimulation, suggesting significant heterogeneity within this compartment.

Finally, to visualize protein and transcriptome data in an intuitive single plot, we adapted One-SENSE, which was originally developed for visualization of mass cytometry data (Cheng et al., 2016). This adaptation allows for effective visualization and identification of cellular phenotypes that differ either by transcript or by protein. Overall, we provide a methodological toolset for generating high throughput multi-omic single cell data with a focus on maximizing target transcript sensitivity at minimal read depth and an analytical tool to visualize these protein and transcript datasets.

## Results

### Comparison of oligonucleotide-labelled antibody probes to high-dimensional flow cytometry

For our reference data set we obtained peripheral blood mononuclear cells (PBMCs) from three healthy control subjects carrying the HLA-A*02:01 allele, which allowed isolation of EBV-specific CD8^+^ T cells using an EBV-Tetramer reagent (Dunne et al., 2002). To ensure sufficient cell numbers of these rare, antigen-specific T cells, we enriched tetramer-positive T cells by fluorescence-activated cell sorting (FACS). In parallel, we sorted CD45^+^ live leukocytes from PBMCs (Figure 1A). Moreover, to minimize batch effects during subsequent staining with 41 oligo-nucleotide labelled antibodies (Figure 1B), we utilized a multiplexing protocol using barcoded cell-hashing antibodies (Stoeckius et al., 2018). All samples were processed simultaneously using the Rhapsody platform, a nano-well based cartridge system (Fan et al., 2015) for single-cell RNA sequencing with a targeted approach focusing on 490 immune-relevant transcripts (all targets are listed in Suppl Table 1). Following quality control and removal of multiplets, we recovered 27,258 cells from the sequencing data, which were evenly distributed across the three different donors.

**Figure 1:**
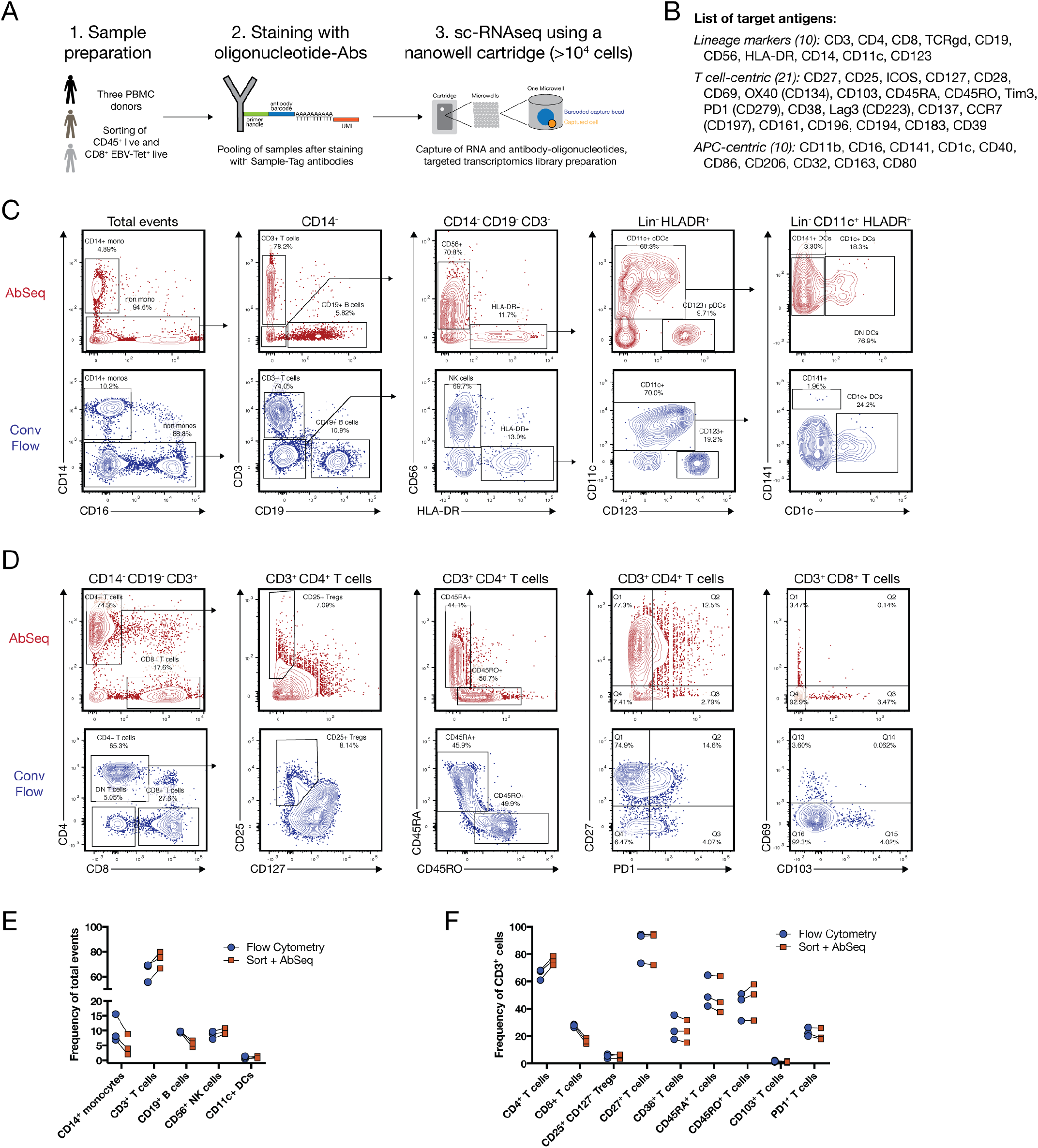
Comparison of oligo-nucleotide antibody probes to high-dimensional flow cytometry. (A) Schematic graph describing the workflow of the experiment. PBMC samples from three donors were split in half, with one aliquot used for the multi-omic workflow, and one aliquot used for flow cytometry phenotyping using two 30-parameter panels. (B) Overview of antibody targets used in both the multi-omic and conventional flow cytometry experiment. (C) Manual gating of main immune subsets using the combined AbSeq data set (upper panel, red) and concatenated and down-sampled events (27,000 cells) from the conventional (conv) flow cytometry data set (lower panel, blue). (D) Manual gating of various T cell markers using the combined AbSeq data set (upper panel, red) and concatenated, down-sampled events from the cytometry data set (lower panel, blue). (E) Quantification of main immune subsets in the AbSeq and flow cytometry data set across the three different donors. (F) Quantification of main T cell populations and selected phenotyping markers in the AbSeq and flow cytometry data set across the three different donors.

First, we wanted to assess whether the surface protein phenotypes as defined by sequencing match known biology. For this, we designed two optimized 30-parameter immunophenotyping panels (adapted from (Mair and Prlic, 2018)) covering the same 41 protein targets in an overlapping fashion. We used these panels to stain whole unsorted PBMC samples from the same 3 donors, down-sampled the cytometry data to 27,000 cells and used biaxial gating to identify the main immune lineages of the myeloid compartment (Figure 1C) as well as the lymphoid compartment (Figure 1D). All populations were present at comparable frequencies in the two different data sets (Figure 1E and Figure 1F), with myeloid cells showing slightly lower abundance due to the sorting procedure required to enrich EBV-Tetramer^+^ cells as well as CD45^+^ live cells. Of note, even low-abundance cell populations such as CD1c^+^ conventional dendritic cells (cDCs) and crosspresenting CD141^+^ cDCs were clearly identified by their surface protein phenotype. Furthermore, the oligonucleotide-labelled antibodies allowed to discriminate the CD45 splice variants CD45RO and CD45RA, which cannot be distinguished by 3’ transcriptomic analysis alone.

However, for the anti-TCRγδ reagent we used, discordant patterns were observed when comparing the expression within CD3^+^ T cells to conventional flow cytometry (Supplementary Figure 1A). This was not immediately evident when visualizing the data on a heatmap (Supplementary Figure 1B), emphasizing the need for careful reagent validation for sequencing-based protein measurements. Thus, we did not analyze γδ T cells separately for the rest of our study. Furthermore, the CCR7 reagent delivered sub-optimal but usable resolution (data not shown).

### Targeted transcriptomics faithfully captures cellular heterogeneity similar to whole transcriptome approaches at lower read depths

Next, we wanted to assess how well a targeted transcriptomics approach can identify immune cell heterogeneity compared to a commonly used whole transcriptome (WTA) pipeline. For this, we used a single donor and compared the resulting populations after graph-based-clustering of the transcript data using the R package Seurat implementation of PhenoGraph at standard resolution settings (Butler et al., 2018; Levine et al., 2015) (Figure 2A and Suppl. Figure 2A and 2B). For visualization, we used uniform manifold approximation and projection (UMAP), a dimensionality reduction approach that has recently been adopted for single-cell data (Becht et al., 2018; McInnes et al., 2018). Overall, the targeted transcriptomic approach utilizing 490 genes revealed similar or even improved resolution of known immune subsets in the peripheral blood. In particular, CD4^+^ T cells and CD8^+^ T cells separated well, and we observed regulatory T cells (Tregs) expressing FOXP3 and CTLA4 as a separate cluster (Figure 2B). For verification of this Treg cluster, we utilized the corresponding protein signature, which showed high expression of CD25, and low expression of CD127 (Figure 2C). Next, we compared the gene expression for four phenotypically similar clusters in the WTA and the targeted transcriptomics data set, showing very similar patterns for the top differentially expressed genes (Suppl. Figure 2B). To obtain a relative measure of detection efficiency, we calculated the average number of transcripts per cell both for the targeted transcriptomics as well as the WTA data set from the same donor. Around 75% of the assayed genes showed equal or slightly superior detection efficiencies (Figure 2D), suggesting that targeted transcriptomics can deliver valuable information at relatively low sequencing cost (i.e. approximately 2500 reads/cell).

**Figure 2:**
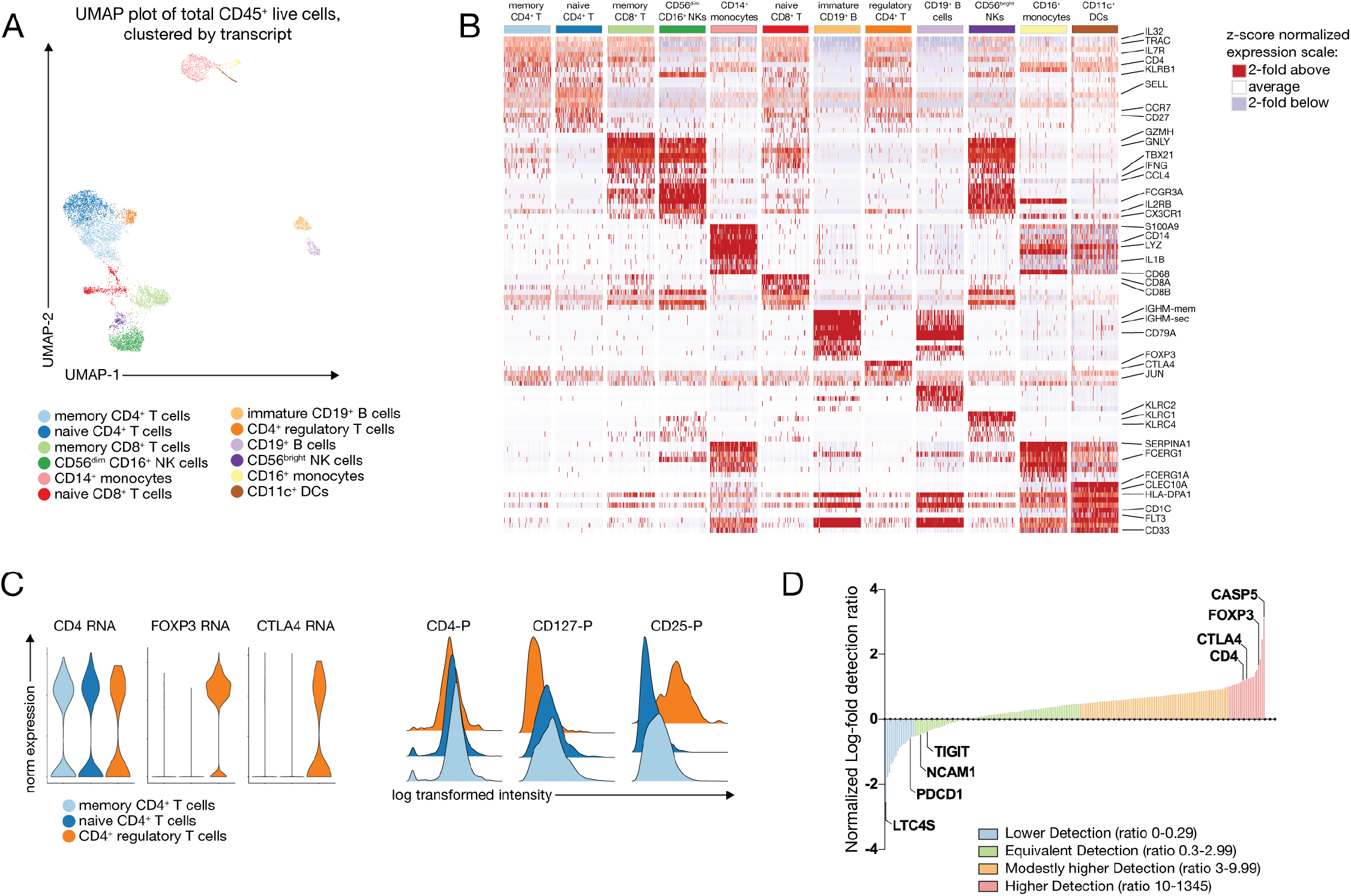
Targeted transcriptomics faithfully captures cellular heterogeneity in peripheral blood mononuclear cells. (A) Graph-based clustering of the transcript data from one representative donor is shown on a UMAP (uniform manifold approximation projection) plot. Clusters have been annotated by expression of key lineage genes. (B) The top 10-differentially expressed genes for each cluster were identified using the Seurat implementation of MAST (model-based analysis of single-cell transcriptomes) and visualized on a heatmap after z-score normalization. Cluster names are shown in the same color scheme as in (A). (C) Expression of the indicated transcripts and proteins on the three different CD4^+^ T cell clusters, highlighting the CD25^+^ CD127^low^ Treg cluster. (D) Relative detection ratio of all detected transcripts relative to a whole transcriptome data set from the same donor. Genes are manually assigned into four different groups according to their relative detection ratio.

Finally, to directly assess the effect of different read-depths on resolution of protein and transcript signals, we analyzed a different donor to a total of approximately 27,000 reads/cell (approximately 18,000 reads/cell for the antibody library, 9,000 reads/cell for the transcript library) and subsampled the number of reads used during processing of the raw data to 20% (approximately 4000 reads/cell for antibody library, 2000 reads/cell for transcript library) and 10%. Visualization of the resulting clusters on a UMAP plot as well as the top-differentially expressed genes on a heatmap revealed no major differences between using 100% or 20% of the reads (Supplementary Figure 2C). For the protein signal, the same was observed, while using only 10% of the reads resulted in noticeable loss of signal intensities (Supplementary Figure 2D). Overall, we conclude that using at least 2000 reads/cell for the transcript portion of the library and at least 200 reads/antibody/cell for the antibody portion of the antibody library delivers sufficient resolution.

### Multi-omic analysis identifies canonical memory T cell populations and allows the study of rare-antigen specific CD8^+^ T cells

To test the value of multi-omic single cell analysis on a specific subset of the immune compartment, we performed an in-depth analysis of the CD8^+^ T cell compartment. First, we visualized protein and RNA data collected from total CD45^+^ live cells from PBMCs from three patents on separate UMAP plots (Fig 1A). We found that cells from different donors comingled and separated by cell type rather than by donor, suggesting that batch effect across donors was minimal (Figure 3A). Of note, protein information overlayed on the transcript-generated UMAP plot allowed accurate identification of all main immune clusters (Figure 3B), which is not necessarily the case when using transcript information for the corresponding lineage markers. This is exemplified by biaxial plots showing protein signal on the y-axis and transcript signal on the x-axis (Figure 3C): While for CD8A, transcript and protein are co-expressed in most cells, only half of the CD4-protein^+^ (throughout the manuscript abbreviated as CD4-P) cells contained detectable CD4-transcript. In turn, there were other molecules of interest where the inverse was observed: CD69-RNA (plotted on the x-axis) was detected across a large number of T cells, but as expected only few T cells in the peripheral blood express CD69 protein (CD69-P, plotted on y-axis) on their surface. For CD27, we observed a higher correlation between transcript and protein (Figure 3C). Overall, these observations emphasize the importance of parallel measurement of protein and transcript to faithfully study T cell biology.

**Figure 3:**
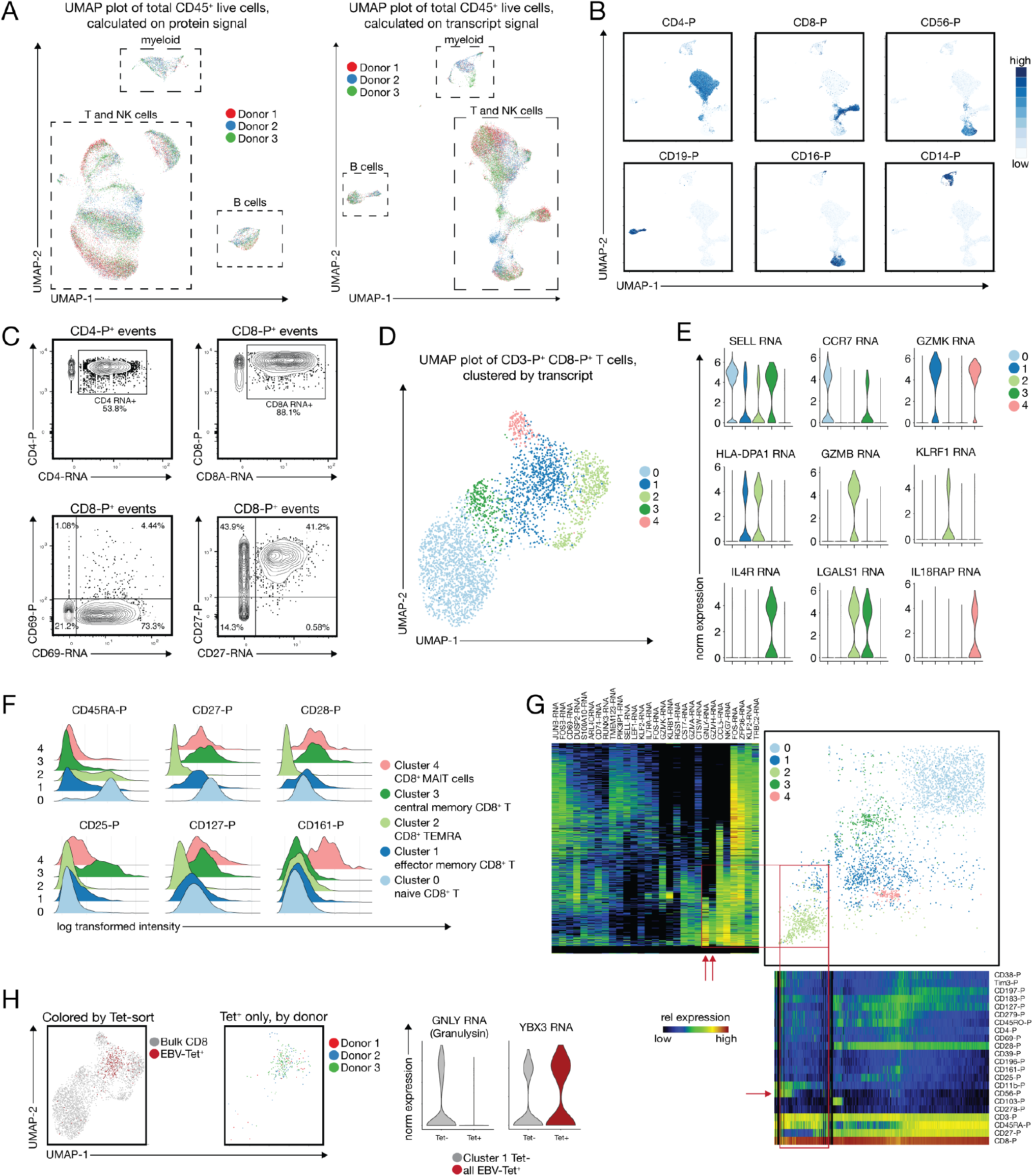
Multi-omic targeted transcriptomics identifies canonical memory T cell populations and allows the study of rare-antigen specific CD8^+^ T cells. (A) UMAP plots calculated on protein (left) or transcript (right) show that there is no batch effect across the three donors analyzed. (B) Example UMAP plots (calculated on transcript) representing the expression of the main immune lineage protein markers which allow the unequivocal identification of CD4^+^ and CD8^+^ T cells, CD19^+^ B cells, and CD14^+^ as well as CD16^+^ myeloid cells. (C) Example plots showing the poor correlation of transcript and protein levels for CD4 and CD69, and good correlation for CD8 and CD27. Protein signal is plotted on the y-axis, transcript on the x-axis. (D) UMAP plot and graph-based clustering of the CD3^+^ CD8^+^CD4^−^ T cell compartment, revealing 5 distinct populations. (E) Examples of top differentially expressed genes identified by MAST for each of the 5 clusters highlighted in (D). (F) Protein signatures of the 5 clusters identified canonical naive and memory CD8^+^ T cell subsets, including mucosal associated invariant T cells (MAIT cells). (G) One-SENSE plot depicting protein expression heatmap along the x-axis, and transcript expression heatmap of the top differentially expressed genes along the y-axis. (H) Identification of EBV-specific CD8^+^ T cells relative to all CD8^+^ T cells, and expression pattern of two differentially expressed genes between Tetramer-positive cells and Tetramer negative cells in the effector memory cluster 1.

Next, we continued our analysis of CD3^+^CD4^−^CD8^+^ T cells as defined by surface protein expression using SCAMP (Selected Clustering Annotated using Modes of Projections) (Greene et al., 2018). Unbiased graph-based clustering using transcript information suggested the presence of 5 distinct cellular clusters (Figure 3D). Visualization of the top differentially expressed genes such as SELL (encoding CD62L), CCR7 and GZMB suggested that these 5 clusters reflect canonical naïve and memory T cell populations (Sallusto et al., 1999) (Figure 3E). Additionally, our data allowed identification of CD8^+^ mucosal associated invariant T (MAIT) cells, which express high levels of IL18RAP and TNF (Slichter et al., 2016) (Mori et al., 2016). We confirmed the resemblance of these populations by surface protein expression (Figure 3F), with central memory CD8^+^ T cells expressing low levels of CD45RA-protein, and high CD27- and CD28-protein (Sallusto et al., 2004) (Hamann et al., 1997). Of note, the splice variants CD45RO and CD45RA cannot be distinguished by analyzing transcript alone, highlighting the added value of combined protein and transcript analysis.

To visualize the correspondence between transcript and protein expression in the multi-omic data set, we adopted One-SENSE, which has originally been developed for biologically meaningful visualization of mass cytometry data (Cheng et al., 2016). For this, we mapped cells separately by proteins and transcripts each on to a single UMAP dimension, similar to a recently published 1D t-stochastic neighbor embedding (t-SNE) representation for sc-RNA sequencing data (Linderman et al., 2019). The combined plot shows the overall distribution of protein expression profiles in the x-axis and the top-differentially expressed gene profiles on the y-axis. Aligned heatmaps that represent median expression with bins of cells are provided to annotate the one-dimensional UMAP protein and gene expression profiles. This approach allows easy identification of cellular clusters that are similar by transcript, but separated by protein, and vice versa (Figure 3G). One example for this is highlighted in Figure 3G (red box and arrow), where cluster 2 (light green, containing TEMRA cells) is relatively homogenous by transcript, but can be separated by CD56 protein expression, probably marking some NKT cells. In turn, a fraction of cells between cluster 1 (dark blue, effector memory CD8^+^ T cells) and 2 (green, TEMRA) shares the same protein signature, but can be distinguished by GNLY and GZMH expression (Fig. 3G, red box and arrows). Varying degrees of concordance and ability to discriminate cellular subsets from gene and protein expression profiles can be seen across this plot.

To determine if targeted transcriptomics is amenable for studying rare antigen-specific T cell populations, we analyzed CD8^+^ T cells recognizing an EBV-epitope (Dunne et al., 2002). Visualization on the UMAP plot revealed remarkable similarity of EBV-specific T cells across all three donors (Figure 3H). As expected, most of the cells grouped within the effector memory CD8^+^ T cell cluster. However, relative to the EBV-nonspecific memory T cell cluster the EBV-Tet^+^ T cells showed a significant downregulation of the effector molecule Granulysin, and an upregulation of YBX3, an RNA binding protein whose function has not been defined in T cells, but has recently been shown to be a critical regulator for the stability of specific mRNAs (Cooke et al., 2019).

Overall, this data show that combining targeted transcriptomics and protein phenotyping by sequencing is a valuable approach for studying T cell subsets and could be used a resource-efficient tool for studying T cell responses in human disease.

### Short-term stimulation of T and NK cells reveals chemokine heterogeneity and a disconnect with the early activation marker CD69

Cytokines and chemokines are the quintessential effector molecules of T cells, and the existence of specific T cell subsets that are poised for the production of certain cytokines has been the subject of intense research over the past decade (van den Broek et al., 2018; Zhou et al., 2009). To test whether multi-omic single-cell analysis can provide additional insight, we purified pan T cells together with NK cells and stimulated them for one hour with Phorbol-Myristate-Acetate (PMA) and Ionomycin. We probed early transcriptional changes with a T cell centric targeted transcriptomic approach covering 259 genes. Transcripts encoding for IFNG, FASL and ICOS exhibited robust upregulation in the stimulated versus unstimulated sample (Figure 4A), as was the case for CD69, a commonly utilized protein marker for early T cell activation (Figure 4B). Of note, when we analyzed cytokine expression relative to the surface protein expression of CD69, we observed that both IFNg as well as TNF transcript was primarily expressed in CD69-transcript positive, but CD69-protein negative cells, suggesting that during very early stages of activation, CD69 protein might not be an ideal marker for T cell activation. However, FOSB, part of the transcription factor AP-1, was co-expressed with CD69-protein (Figure 4B), suggesting a close relationship of FOSB and CD69 expression.

**Figure 4:**
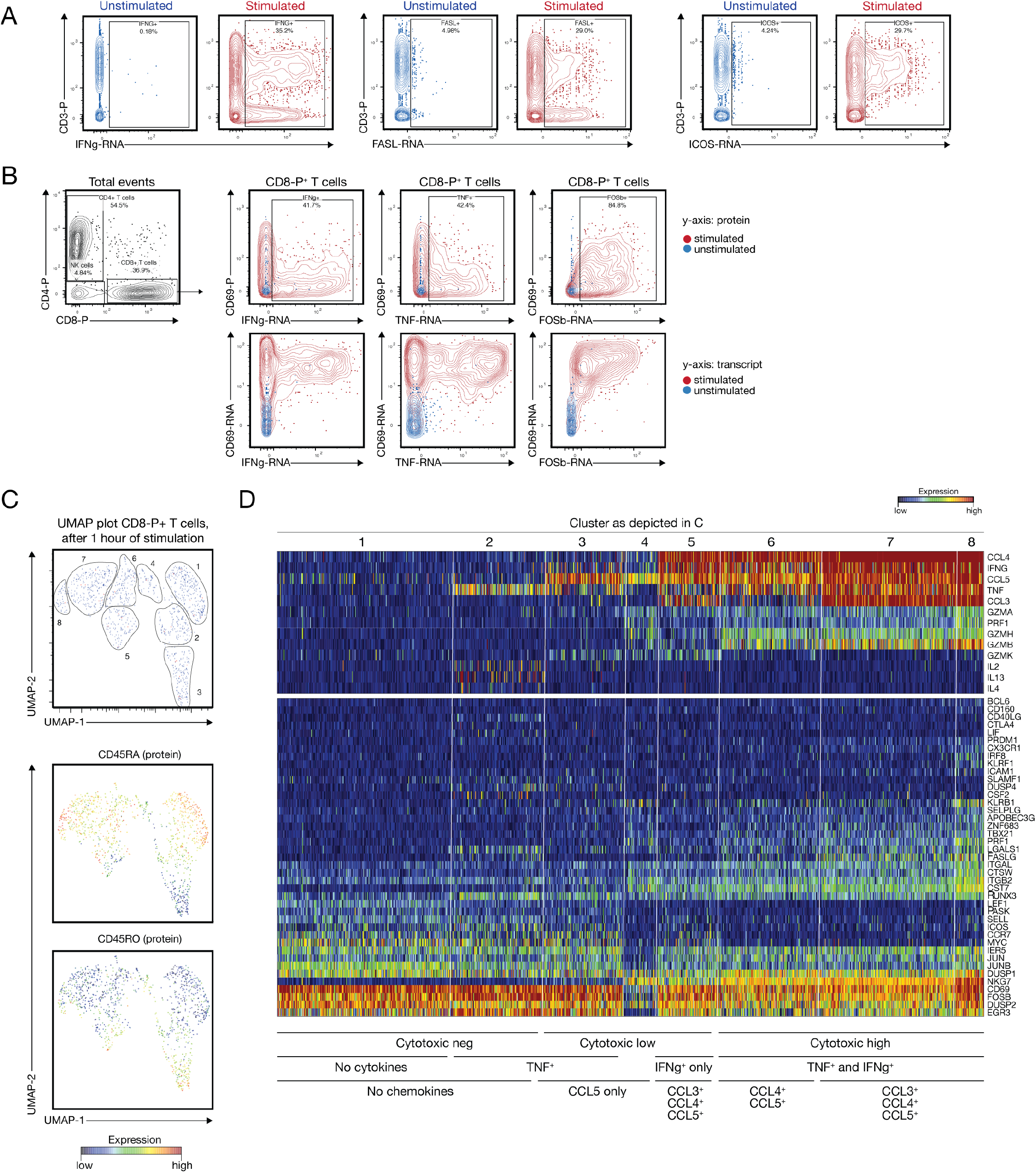
Multi-omic analysis of the T and NK cell compartment 1 hour after stimulation. (A) Representative plots showing the upregulation of selected effector transcripts such as IFNG, FASL and ICOS after stimulation (red) relative to unstimulated cells (blue). (B) Disconnect between surface protein expression of the early activation marker CD69 and IFNG and TNF transcript within CD8-protein^+^ T cells. Blue overlay indicated unstimulated cells, red stimulated cells. (C) UMAP plot of CD8-protein^+^ T cells with manually identified clusters, and CD45RA and CD45RO protein expression. (D) Heatmap showing the expression of key effector transcripts within the clusters identified in (C).

We focused our further analysis on CD8^+^ T cells only, though our data set also contains information on NK cells. Projection on a UMAP plot showed 8 discernable clusters that were selected manually. Protein expression patterns for CD45RA and CD45RO highlight the naïve and the memory T cells within this plot (Figure 4C). A heatmap visualization of the most highly expressed transcripts show that these clusters are defined by differential expression of CCL3, CCL4, IFNG, TNF, and various granzymes (Figure 4D). Overall, this analysis reveals considerable functional diversity within the CD8^+^ T cell compartment that is detectable as early as one hour after stimulation.

### Multi-omic analysis of the peripheral myeloid compartment reveals inflammatory subsets not captured by surface protein phenotype

Next, we wanted to assess whether the targeted transcriptomics approach can also be used for other immune populations that are not as well studied as T cells. During the past decade it has become evident that the myeloid cell compartment is complex in terms of cellular heterogeneity (Guilliams et al., 2014; See et al., 2017; Villani et al., 2017), and that commonly used bone-marrow derived differentiation protocols do not faithfully capture the phenotype of myeloid cells *in vivo* (Guilliams and Malissen, 2016; Helft et al., 2015). Thus, we tested how well targeted transcriptomics could dissect the heterogeneity of the peripheral myeloid compartment. Unbiased clustering using transcript suggested the presence of 5 different populations (Figure 5A), with clear separation for CD14 and CD16 protein expression (Figure 5B). As expected, visualization of the top differentially expressed genes (Figure 5C) as well as key surface proteins (Figure 5D) mapped these clusters to CD123^+^ plasmacytoid dendritic cells (pDCs), CD1c^+^ conventional DCs (cDC2s), CD16^+^ monocytes and CD14^+^ monocytes. We used One-SENSE to further explore the relationship between cluster 0 and 1, revealing that these two populations were very similar in terms of surface protein profile (CD14^+^CD16^−^), but separated by a specific set of transcripts encoding for pro-inflammatory cytokines and chemokines (Figure 5E). We confirmed that these transcripts were part of differentially expressed genes as identified by MAST (Finak et al., 2015), with higher expression in cluster 1 of CXCL3 and CCL4 (also known as MIP-1b, a chemoattractant for natural killer cells) (Figure 5E). Thus, combining protein and transcriptome data allowed us to observe multiple functional subsets within the peripheral CD14^+^ myeloid population which were not apparent by surface marker expression alone. In summary, this data highlights that targeted transcriptomics can be used for exploratory studies of different immune compartments.

**Figure 5:**
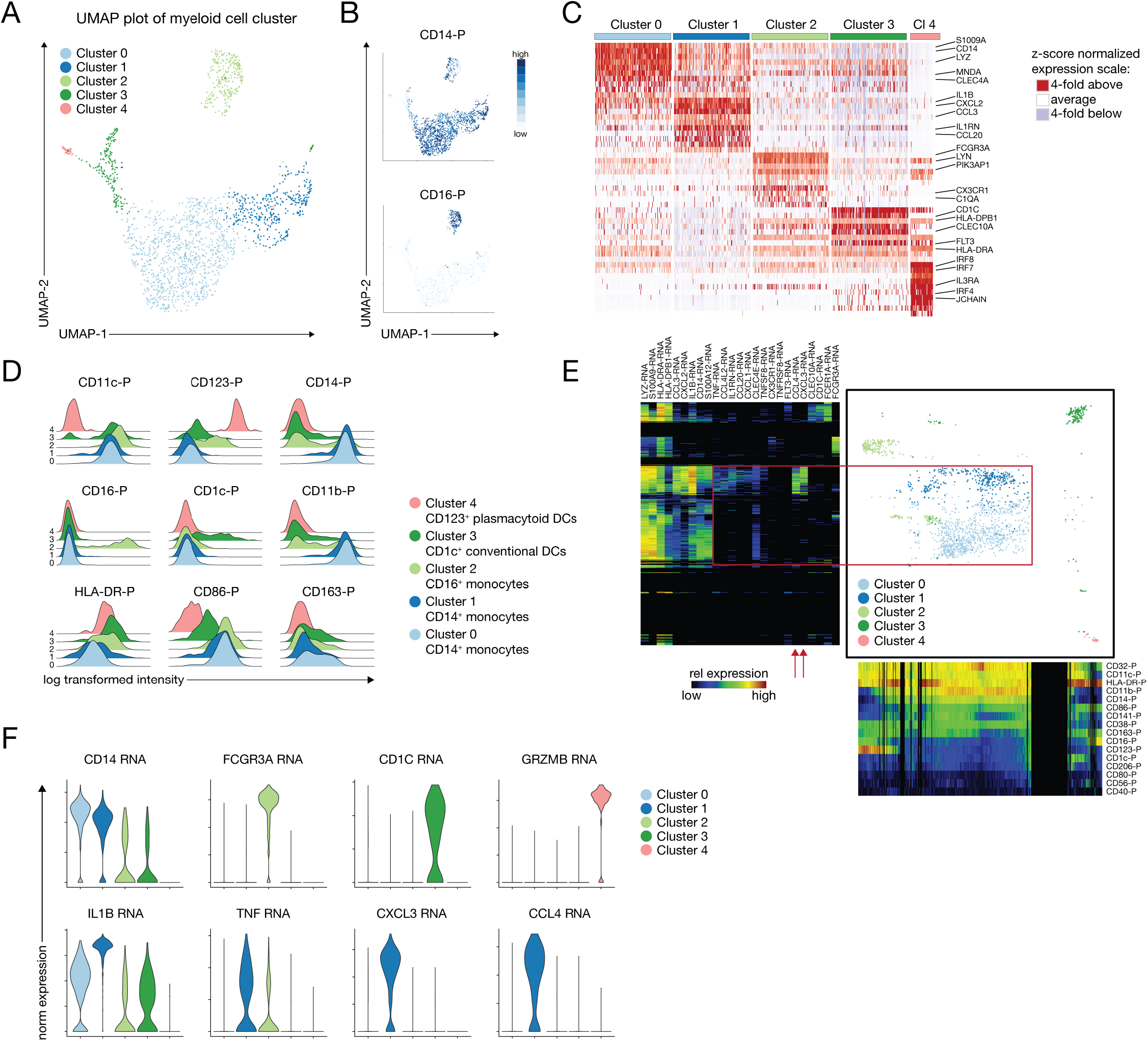
Combined protein and transcript phenotyping of the peripheral myeloid compartment reveals inflammatory subsets not captured by surface protein phenotype. (A) UMAP plot and graph-based clustering of the peripheral non T/non NK/non B cell compartment, revealing 5 distinct populations. (B) Heatmap overlay of CD14^−^ and CD16-protein expression. (C) Heatmap of the top differentially expressed genes identified by MAST for each of the 5 clusters highlighted in (A). (D) Protein signatures of the 5 clusters identifies canonical CD123^+^ plasmacytoid DCs, CD1c^+^ conventional DCs and CD16^+^ monocytes, but two of the clusters mapping to CD14^+^ monocytes. (E) One-SENSE plot depicting protein expression heatmap along the x-axis, and transcript expression heatmap of some of the top differentially expressed genes along the y-axis. Red box and arrrows are highlighting the differentially expressed genes between cluster 0 and 1. (F) Violin plots showing key genes of the respective myeloid population (upper panel) and differentially expressed genes between cluster 0 and 1, suggesting the presence of an inflammatory subpopulation within CD14^+^ CD16^−^ monocytes that expresses high levels of IL1B, TNF, CXCL3 and CCL4.

## Discussion

Current efforts in the field of single cell analysis focus on the integrative measurement of multiple modalities per cell. Ultimately, being able to analyze genome accessibility status, transcript, regulatory RNAs and protein expression all together would allow a holistic understanding of cellular function, but this has not been achieved yet (Stuart and Satija, 2019). Arguably one of the most important steps on this trajectory has been the ability to combine protein and transcript measurements by sequencing at the single cell level using high-throughput methods (Peterson et al., 2017; Stoeckius et al., 2017). However, with increased cell numbers, these combined measurements can quickly become resource intensive, mostly due to the high number of sequencing reads that are required per cell. Moreover, to fully leverage the advantage of multi-omic single-cell analysis approaches, it is imperative to collect large cell numbers to adequately represent low-abundance cellular populations such as antigen-specific T cells, or antigen-presenting cells.

The targeted transcriptomic approach that we describe here provides an alternative platform that significantly lowers the number of reads required for sequencing saturation of transcript compared to whole transcriptome (WTA) approaches, but still provides valuable information on up to 499 immune-centric genes. Though this approach sacrifices the unbiased nature of WTA measurements, many immunological applications center on a set of critical immune effector molecules, such as cytokines, chemokines or transcription factors. Also, a targeted approach avoids the significant number of reads used by transcripts encoding ribosomal proteins which are often also captured using a poly-A based whole transcriptome workflow. Furthermore, as shown here, in some cases, targeted analysis can permit higher sensitivity when it comes to detecting relatively low abundance genes. Overall, in many experimental setups it might be beneficial to combine both approaches: first utilize a WTA platform to identify potentially unknown transcripts, and then use a targeted approach (which can be tailored towards gene sets of interest) for profiling larger cell numbers or interrogating cellular responses to specific stimuli. We provide proof-of-concept data that as early as one hour after stimulation CD8^+^ T cells show heterogeneous patterns of chemokine expression. Comprehensive chemokine and cytokine profiling of T cells after a very short stimulus could be very valuable to gain additional insights into their function e.g. in the context of cancer immunotherapy (Nagarsheth et al., 2017).

The decreased number of reads per cell required for targeted transcriptomics makes the approach very suitable for combined profiling of transcript and protein for larger number of cells. Doing so is particularly relevant in the context of T cell biology, where well established T cell subsets, such as memory T cells and regulatory T cells (Tregs) up to date have been difficult to resolve in some droplet-based sc-RNAseq studies solely on the basis of transcript (Zheng et al., 2017). This has been attributed to the fact that lymphocytes contain a relatively low amount of mRNA, which in combination with the inherent drop-out rate of sc-RNAseq protocols fails to detect some low abundance transcripts that are defining these cellular subsets (Stuart and Satija, 2019). This issue can be alleviated by measuring surface protein markers such as the splice variants CD45RA and CD45RO, which have been well studied in the context of naïve and memory T cells, or the IL-2 receptor alpha chain (CD25) and IL-7 receptor (CD127) for the distinction of Tregs. In addition, parallel measurement of surface protein phenotypes allows to link novel cellular clusters (that are defined solely by transcript) with a large body of literature that used to define cells by surface protein phenotype only. Finally, the combined measurement approach can be useful to identify targets with a significant disconnect between transcript and protein expression such as CD69, probably indicative of active post-transcriptional modifications.

Of note, the development of novel technologies can sometimes outpace our ability to validate platforms and reagents. Given that typical single cell sequencing experiments require complex pre-processing steps and are often visualized using dimensionality reduction techniques such as UMAP or t-SNE, there is a disconnect between the actual raw data and the interpretation of final heatmaps. While this might be less of a problem for transcript counts, antibody-based probes require careful validation. Here, we have used high-dimensional cytometry, highlighting that not all reagents, even if the same antibody clone is used, perform equally well in a multi-omic sequencing experiment relative to conventional cytometry. Thus, with the more widespread adoption of sequencing-based protein measurements, we argue that reagents need to be carefully tested, preferably with parallel deposition in public databases.

Ultimately, to advance our understanding of biology the field relies on innovative approaches to analyze and visualize complex high-dimensional data (Butler et al., 2018; Cao et al., 2019; Stuart and Satija, 2019). Due to the different expression scales this presents a challenge for combined protein-transcript data sets. To alleviate this problem, we have adopted an analysis approach successfully used for high-dimensional cytometry data, one-dimensional soli expression by nonlinear stochastic embedding (One-SENSE) (Cheng et al., 2016). By visualizing the top-differentially expressed genes in one dimension relative to the measured protein phenotypes this method allows to easily dissect cells that are similar in transcript, but different in surface phenotype, and vice versa. This will be a useful tool for biologists to explore future multi-omic data sets to extract biological meaning from these complex multi-dimensional data.

## Acknowledgements

We would like to thank the HIV Vaccine Trial Network (HVTN) for providing samples and for access to their flow cytometry instrumentation (in particular Dr. Stephen de Rosa), the Flow Cytometry Shared Resources Core of the FHCRC (in particular Andrew Berger), and the Genomics Shared Resources Core of the FHCRC for sequencing. We thank members of the Newell and Prlic labs for critical discussion. This work was supported by NIH grants R01AI123323 (to M.P.) and 5U19AI128914 (to R.G.). E.W.N., Y.S., and T.B. were supported by Fred Hutchinson Cancer Research Center New Development funding and by the Andy Hill Research Endowment.

## Author contributions

F.M. and J.R.E. designed and performed experiments, analyzed data and wrote the manuscript. V.V. analyzed data and provided critical input. Y.S. performed experiments. T.B. analyzed data. A.J.T. and J.M. provided critical input. E.W.N., R.G. and M.P. designed the study, analyzed data and co-wrote the manuscript.

## Conflict of interest

A.J.T. and J.M. are employees of BD Biosciences (manuscript approval by BD Biosciences was not required and BD Biosciences had no influence in regard to data analysis, data interpretation and discussion). R.G. has received support from Juno Therapeutics and Janssen Pharma, has consulted for Takeda Vaccines, Juno Therapeutics and Infotech Soft, and has ownership interest in CellSpace Bio.

---

**Supplementary figure 1:**
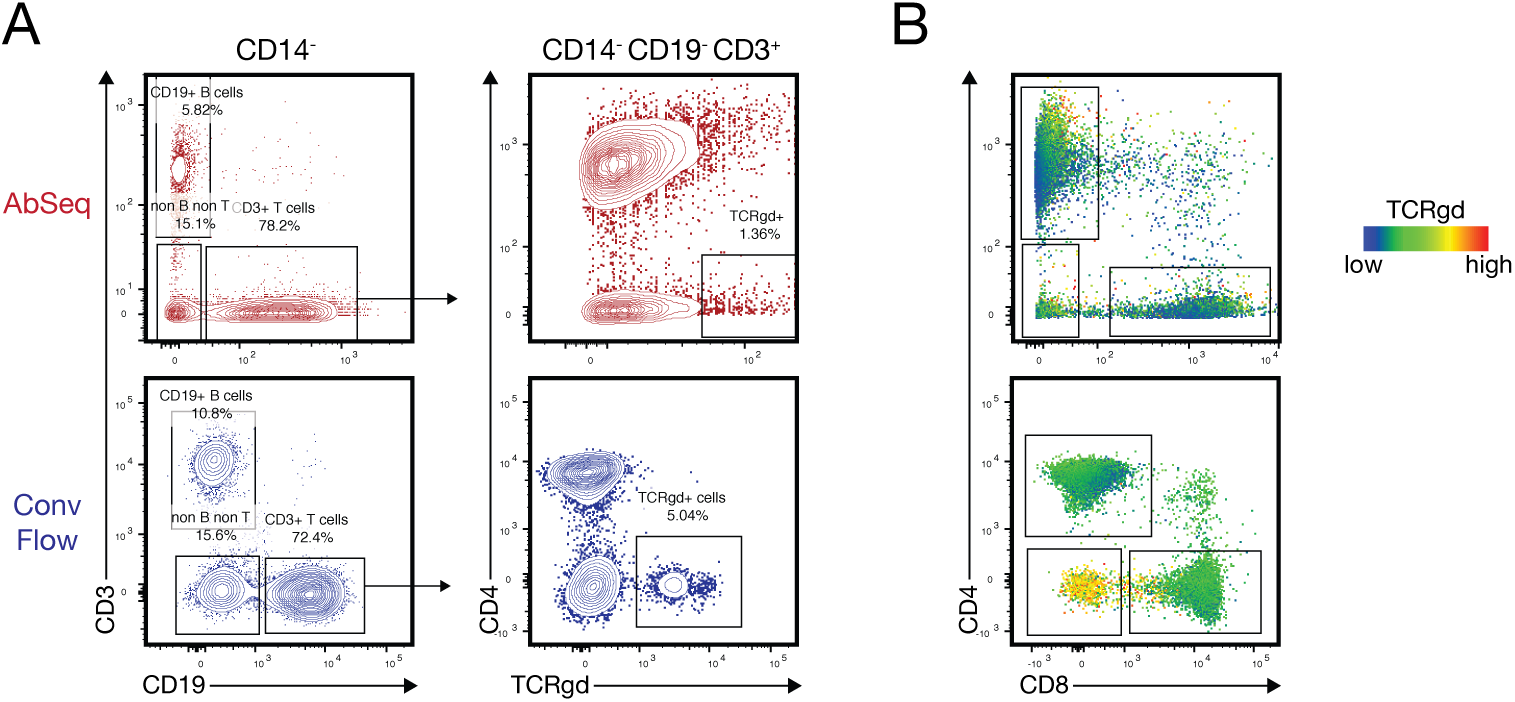
Example for a poorly performing reagent. (A) Manual gating of main immune subsets using the combined AbSeq data set (upper panel, red) and concatenated and down-sampled events from the flow cytometry data set (lower panel, blue), highlighting the population of γδ T cells. (B) Heatmap overlay of the TCRgd signal on a CD4 vs CD8 plot for the AbSeq data set (upper panel) and flow cytometry data set (lower panel).

**Supplementary figure 2:**
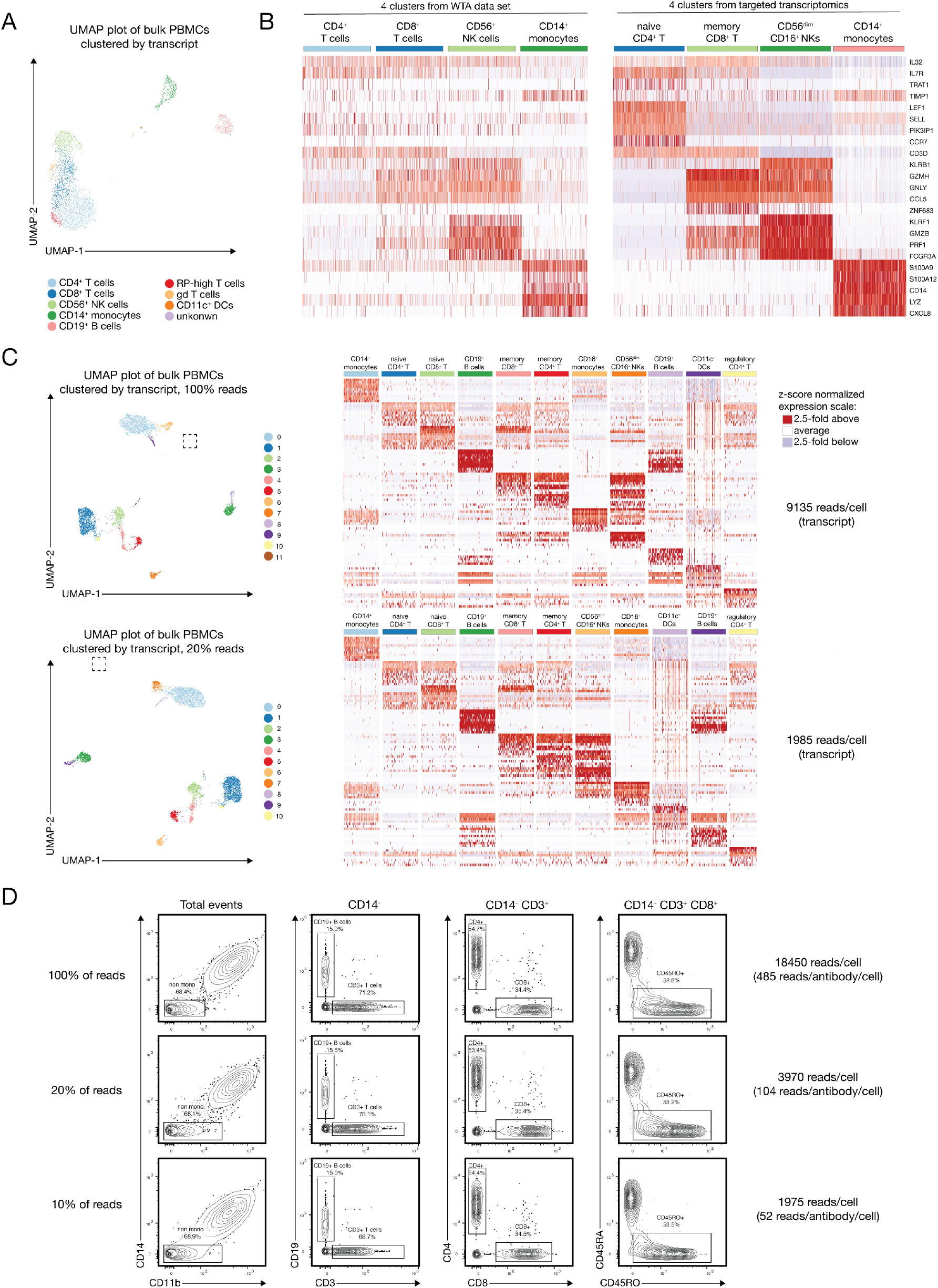
Comparison of targeted transcriptomics to whole transcriptome data (WTA) and assessment of required sequencing depth. (A) Graph-based clustering of WTA data obtained from the same donor as in main Figure 2. (B) Four of the clusters that matched most closely in terms of their expression pattern were selected from both experiments and plotted using the top differentially expressed genes obtained from the targeted transcriptomics approach. Heatmap represents relative expression after z-score normalization. Left plot shows WTA data, right plot shows targeted transcriptomic (cells obtained from the same donor). (C) 5,400 cells from a different donor were sequenced at a total depth of approximately 30,000 reads/cell. Upper panel depicts UMAP plot after graph-based clustering and a heatmap of the top differentially expressed genes (z-score normalized expression) at full read depth, lower panel using only 20% of the reads. Read depth per cell for the transcript library is indicated on the right). Squared box on the UMAP plot indicates one cluster that is separated as cluster 11 at full read depth, but pooled with cluster 8 at lower read-depth (D) Protein signals at the indicated read depths.

## STAR methods

### Cells

Peripheral blood mononuclear cells (PBMCs) were obtained as cryopreserved samples from healthy controls (Seattle Area Control Cohort) via the HIV Vaccine Trial network (HVTN). Vials with cryopreserved cells were thawed at 37°C until a tiny ice crystal was left in the tube, and then carefully diluted in 1mL of pre-warmed RPMI with 10% FBS and transferred to a new tube. An additional 13 mL of pre-warmed RPMI with 10% FBS were added drop by drop, followed by centrifugation for 5 minutes at 400g and resuspension in 1 mL of RPMI.

### Flow Cytometry and Cell sorting

For flow cytometric analysis good practices were followed as outlined in the guidelines for use of flow cytometry (Cossarizza et al., 2017). Following thawing, PBMCs were incubated with Fc-blocking reagent (BioLegend Trustain FcX, #422302) and fixable UV Blue Live/Dead reagent (ThermoFisher, #L34961) in PBS for 15 minutes at room temperature. If required, cells were stained with an EBV-Tetramer reagent (peptide YVLDHLIVV; Fred Hutch Immune Monitoring Core) diluted in FACS buffer (PBS with 2% FBS, Nucleus Biologics) for 30 minutes at room temperature, followed by two washes. After this, cells were incubated for 20 minutes at room temperature with antibody master mix freshly prepared in Brilliant staining buffer (BD Bioscience, # 563794), followed by two washes. All antibodies were titrated and used at optimal dilution, and staining procedures were performed in 96-well round-bottom plates. Stained cells were fixed with 4% PFA for 20 minutes at room temperature, washed, resuspended in FACS buffer and stored at 4°C in the dark until acquisition.

All samples were acquired using a FACSymphony A5 (BD Biosciences), equipped with 30 detectors and 355nm, 405nm, 488nm, 532nm and 628nm lasers and FACSDiva (BD Biosciences). Detector gains were optimized using a modified voltage titration approach (Perfetto et al., 2012) and standardized from day to day using 6-peak Ultra Rainbow Beads (Spherotec, # URCP-38-2K). Single-stained controls were prepared with every experiment using antibody capture beads diluted in FACS buffer (BD Biosciences anti-mouse, #552843 and anti-rat, #552844). After acquisition, data was exported in FCS 3.1 format and analyzed using FlowJo (version 10.5.x, BD Biosciences). Doublets were excluded by FSC-A vs FSC-H gating. For some of the plots, the number of acquired cells was down-sampled using the appropriate FlowJo plugin to match the number of cells analyzed by AbSeq.

All cell sorting was performed on a FACSAria III (BD Biosciences), equipped with 20 detectors and 405nm, 488nm, 532nm and 628nm lasers. For all sorts, an 85 μm nozzle operated at 45 psi sheath pressure was used. Cells were sorted into chilled Eppendorf tubes containing 500 μL of RPMI, washed once in PBS and immediately used for subsequent processing.

### Targeted Transcriptome and protein single-cell library preparation and Sequencing

CD45^+^ live PBMCs and EBV-tetramer^+^ CD8^+^ T cells were sequentially labeled using Single Cell Labelling with the BD Single-Cell Multiplexing Kit and BD AbSeq Ab-Oligos reagents strictly following the manufacturers protocol (BD Biosciences). Briefly, cells from each donor or subtype of cells (after sorting) were labelled with sample tags (Stoeckius et al., 2018). Each sample was washed twice with FACS buffer before pooling all samples together. Pooled samples were washed one more time and then stained with AbSeq Ab-Oligos (BD Biosciences). The pooled sample was then washed twice, counted and resuspended in cold BD Sample Buffer (BD Biosciences) to achieve approximately 20,000 cells in 620 μl. Single cells from the pooled sample were isolated using Single Cell Capture and cDNA Synthesis with the BD Rhapsody Express Single-Cell Analysis System following the manufacturers protocol (BD Biosciences). After priming the nanowell cartridges, the pooled sample was loaded onto two BD Rhapsody cartridges and incubated at room temperature. Cell Capture Beads (BD Biosciences) were prepared and then loaded onto the cartridge and incubated prior to shaking at 1,000rpm at room temperature for 15 seconds on a ThermoMixer C (Eppendorf). According to the manufacturers protocol, cartridges were washed, cells were lysed, and Cell Capture Beads were retrieved and washed prior to performing reverse transcription and treatment with Exonuclease I. cDNA Libraries were prepared using mRNA Targeted, Sample Tag, and BD AbSeq Library Preparation with the BD Rhapsody Targeted mRNA and AbSeq Amplification and BD Single-Cell Multiplexing Kits and protocol (BD Biosciences). In brief, cDNA underwent targeted amplification using the Human Immune Response Panel primers and a custom supplemental panel (all targets are listed in Supplementary Table 1) via PCR (10 cycles). PCR products were purified, and mRNA PCR products were separated from sample tag and AbSeq products with double-sided size selection using SPRIselect magnetic beads (Beckman Coulter). mRNA and Sample Tag products were further amplified using PCR (10 cycles). PCR products were then purified using SPRIselect magnetic beads. Quality and quantity of PCR products were determined by using an Agilent 2200 TapeStation with High Sensitivity D5000 ScreenTape (Agilent) in the Fred Hutch Genomics Shared Resource laboratory. Targeted mRNA product was diluted to 2.5 ng/μL and sample tag and AbSeq PCR products were diluted to 1 ng/μL to prepare final libraries. Final libraries were indexed using PCR (6 cycles). Index PCR products were purified using SPRIselect magnetic beads. Quality of final libraries was assessed by using Agilent 2200 TapeStation with High Sensitivity D5000 ScreenTape and quantified using a Qubit Fluorometer using the Qubit dsDNA HS Kit (ThermoFisher). Final libraries were diluted to 2nM and multiplexed for paired-end (150bp) sequencing on a HiSeq 2500 sequencer (Illumina).

### Whole Transcriptome single-cell library preparation and sequencing

cDNA libraries of CD45^+^ Live PBMCs were generated using the Chromium Single Cell 3’ Reagent Kits v2 (10x Genomics) protocol targeting 5,000 cells in two separate wells. Briefly, single cells were isolated into oil emulsion droplets with barcoded gel beads and reverse transcriptase mix. cDNA was generated within these droplets, then the droplets were dissociated. cDNA was purified using DynaBeads MyOne Silane magnetic beads (ThermoFisher). cDNA amplification was performed by PCR (10 cycles) using reagents within the Chromium Single Cell 3’ Reagent Kit v2 (10x Genomics). Amplified cDNA was purified using SPRIselect magnetic beads (Beckman Coulter). cDNA was enzymatically fragmented and size selected prior to library construction. Libraries were constructed by performing end repair, A-tailing, adaptor ligation, and PCR (12 cycles). Quality of the libraries was assessed by using Agilent 2200 TapeStation with High Sensitivity D5000 ScreenTape (Agilent). Quantity of libraries was assessed by performing digital droplet PCR (ddPCR) with Library Quantification Kit for Illumina TruSeq (BioRad). Libraries were diluted to 2nM and paired-end sequencing was performed on a HiSeq 2500 sequencer (Illumina).

### Cell Ranger processing for WTA data

Raw base call (BCL) files were demultiplexed to generate Fastq files using the cellranger mkfastq pipeline within Cell Ranger 2.1.1 (10x Genomics). Targeted transcriptome Fastqs were further analyzed via Seven Bridges (BD Biosciences). Whole transcriptome Fastq files were processed using the standard cellranger pipeline (10x genomics) within Cell Ranger 2.1.1. Briefly, cellranger count performs alignment, filtering, barcode counting, and UMI counting. The cellranger count output was fed into the cellranger aggr pipeline to normalize sequencing depth between samples. The final output of cellranger (molecule per cell matrix) was then analyzed in R using the package Seurat (version 2.3 and 3.0) as described below.

### Seven Bridges processing for targeted transcriptomics data

Targeted transcriptomics Fastq files were processed via the standard Rhapsody analysis pipeline (BD Biosciences) on Seven Bridges (www.sevenbridges.com). First, R1 and R2 reads are filtered for high-quality reads, dropping reads that are too short (less than 64 bases for R2) or have a base quality score of less than 20. Then, R1 reads are annotated to identify cell label sequences and unique molecular identifiers (UMIs), and R2 reads are mapped to the respective reference sequences using Bowtie2. Finally, all valid R1 and R2 reads are combined and annotated to the respective molecules. For all of our analysis, we utilized recursive substation error correction (RSEC) as well as distribution-based error correction (DBEC), which are manufacturer-developed algorithms correcting for PCR and sequencing errors. For determining putative cells (which will contain many more reads than noise cell labels), a filtering algorithm takes the number of DBEC-corrected reads into account, calculating the minimum second derivative along the cumulative reads as the cut-off point. Final expression matrices contain DBEC-adjusted molecule counts in a CSV format. For further analysis, these molecule count tables were read into the R package Seurat (version 2.3 and 3.0) using customized scripts and analyzed as described below.

### Seurat workflow for targeted and WTA data

The R package Seurat (Butler et al., 2018) was utilized for all downstream analysis. For whole transcriptome data, cells that had at least 200 genes (with ≤ 20% being mitochondrial genes) were included in analysis. A natural log normalization using a scale factor of 10,000 was performed across the library for each cell. UMIs and mitochondrial genes (only for WTA data) were linearly scaled to remove these variables as unwanted sources of variation. Dimensionality reduction using UMAP and clustering was performed on a subset of variable genes. For targeted transcriptomics, no gene per cell cutoffs were imposed, data were normalized with the same method. However, when scaling data, UMI was the only regressed variable. Dimensionality reduction using UMAP and clustering was based on either all genes or all proteins. For differential gene expression analysis we utilized the Seurat implementation of MAST (model-based analysis of single-cell transcriptomes) (Finak et al., 2015). For generation of some FCS files the antibody molecule count tables were converted using the R packages premessa and flowCore. FCS-files with antibody molecule count signals were analyzed in FlowJo 10.5.x (BD Biosciences) using either an arcsin transform or biexponential transform. All the scripts used, listing the detailed parameters for each step are available at https://github.com/MairFlo/Targeted_transcriptomics. Raw data will be deposited on the NCBI gene expression Omnibus at https://www.ncbi.nlm.nih.gov/geo/.

### Data processing for One-SENSE and generation of FCS files

CSV files of raw counts were converted to FCS files using a script adapted from https://gist.github.com/yannabraham/c1f9de9b23fb94105ca5. Raw counts were normalized based on total counts per cell, then scaled to a value of 10,000 based on the Seurat normalization algorithm. A natural log transformation was applied to gene expression data, while protein expression data was randomized by adding a random uniform distribution from 0 to 1, followed by transformation with the function arcsinh(x/5). Dimensionality reduction using UMAP was performed separately on all genes and proteins to reduce them to one dimension before plotting. Cells were also split into 500 bins of equivalent width based on one-dimensional UMAP data, then used to generate heatmaps colored by median marker intensity per bin. All scripts used for data processing and plot generation are available at https://github.com/MairFlo/Targeted_transcriptomics.

